# Sex-specific age-related changes in excitatory and inhibitory intra-cortical circuits in mouse primary auditory cortex

**DOI:** 10.1101/2022.06.15.496332

**Authors:** Binghan Xue, Joseph P-Y Kao, Patrick O. Kanold

## Abstract

A common impairment in aging is age-related hearing loss (presbycusis), which manifests as impaired spectrotemporal processing. Aging is accompanied by alteration in normal inhibitory (GABA) neurotransmission and changes in excitatory (NMDA and AMPA) synapses in the auditory cortex (ACtx). However, the circuit mechanisms responsible for age-related auditory dysfunction remain unknown. Here we investigated how auditory cortical microcircuits change with age. We performed laser-scanning photostimulation (LSPS) combined with whole-cell patch clamp recordings from Layer (L) 2/3 cells in primary auditory cortex (A1) in young adult (postnatal day (P) 47-P72) and aged (P543 to P626) male and female CBA/CaJ mice. We found that L2/3 cells in aged male animals display functional hypoconnectivity of both excitatory and inhibitory circuits originating from L4. Compared to cells from young adult mice, cells from aged male mice have fewer inhibitory connections from L4 while female mice show weaker connection strength. These results suggest a sex-specific reduction in excitatory and inhibitory intralaminar cortical circuits in aged mice compared with young adult animals. We speculate that these unbalanced changes in cortical circuits contribute to the functional manifestations of age-related hearing loss in both males and females.

## Introduction

Sensory and cognitive declines with aging are profound and likely involve changes in cortical processing (Nusbaum, 1999; Scialfa, 2002; Jayakody et al., 2018). One common impairment is age-related hearing loss, also known as presbycusis (Brant and Fozard, 1990; Pearson et al., 1995; Matthews et al., 1997; Cruickshanks et al., 1998). Age-related loss of high-frequency hearing has been associated with changes in tonotopic organization and deterioration of temporal processing in the inferior colliculus (IC) and auditory cortex (ACtx) (Willott et al., 1993; Mendelson and Ricketts, 2001; Lee et al., 2002; Mendelson and Lui, 2004; Gourevitch and Edeline, 2011; Engle and Recanzone, 2013; Trujillo et al., 2013; Trujillo and Razak, 2013; Brewton et al., 2016; Recanzone, 2018; Shilling-Scrivo et al., 2021). Moreover, an abnormal tuning bandwidth has been observed in aging rats (Turner et al., 2005). *In vivo* imaging of the primary auditory cortex (A1) in mice of the CBA strain, which retains good peripheral hearing into adulthood (Willott et al., 1988, 1991; Bowen et al., 2020), has shown that aging leads to a reduction in the diversity of tuning curves, a lack of suppressive responses, and increased correlated activity (Shilling-Scrivo et al., 2021). Given that activity correlations are determined by both ascending inputs to L2/3 as well as intra-laminar connections within L2/3 this suggested that intracortical circuits are altered with aging. However, it is unknown if the functional connectivity of microcircuits in A1 changes with aging.

The interplay of glutamatergic excitation and GABAergic inhibition contributes to normal brain function. Disruption of inhibition is thought to be responsible for age-related impairment of suprathreshold signals including auditory signal in noise. (Chao and Knight, 1997; Kok, 1999; Caspary et al., 2008; Stanley et al., 2012; Rozycka and Liguz-Lecznar, 2017; Recanzone, 2018; Shilling-Scrivo et al., 2021). Indeed, aging decreases the density of inhibitory synapses in the prefrontal cortex (Peters et al., 2008). It is also associated with decreased levels of glutamate decarboxylase (GAD) and vesicular GABA transporter (VGAT), hypofunction of NMDA receptors, and changes in ionic conductances in different sensory cortices (De Luca et al., 1990; Chaudhry et al., 1998; Milbrandt et al., 2000; Shi et al., 2004; Stanley and Shetty, 2004; Ling et al., 2005; Burianova et al., 2009; Stanley et al., 2012; Gold and Bajo, 2014; Liguz-Lecznar et al., 2015; Liao et al., 2016; Kumar et al., 2019). Interestingly, administering GABA or GABA agonists facilitates visual function in vision-impaired aged animals (Leventhal et al., 2003). Similarly, GABA-associated changes are observed in the auditory pathway in aged animals. Aging alters GABA receptor composition, the levels of GAD and calcium binding proteins (CBPs) in the IC, thalamus, cochlear nucleus, and ACtx (Milbrandt et al., 1996; Caspary et al., 1999; Ling et al., 2005; Ouda and Syka, 2012; Caspary et al., 2013; Richardson et al., 2013; Brewton et al., 2016; Recanzone, 2018; Richardson et al., 2020; Rogalla and Hildebrandt, 2020). These results suggest that the primary functional deficit in cortical circuits in aging may lie in a hypofunction of inhibitory circuits. At the same time, however, synaptic excitation also changes with aging, since the density of dendritic spines is decreased in aged brains and the structure of the NMDA receptor complex changes with aging (Magnusson et al., 2010; Dickstein et al., 2013; Hickmott and Dinse, 2013; Fetoni et al., 2015). These results suggest that both excitatory and inhibitory circuits may be altered in aged brains.

We thus investigated the changes in the functional microcircuits of A1 in CBA mice with aging. Importantly, while mice of the C57BL/6J strains have peripheral hearing loss in young adulthood (Henry and Chole, 1980; Willott, 1986; Spongr et al., 1997) and show a remapping of ACtx in response to peripheral hearing loss (Willott et al., 1993), CBA mice have good peripheral hearing into old age (Willott et al., 1988, 1991; Frisina et al., 2011). Moreover, in vivo imaging of A1 of CBA mice showed an unchanged distribution of high-frequency responding neurons in A1 at 15-17 months of age (Shilling-Scrivo et al., 2021), suggesting a largely intact peripheral auditory system at this age. This allowed us to investigate the A1 circuit changes with aging. We used laser-scanning photostimulation (LSPS) in combination with whole-cell patch clamp recordings of A1 L2/3 cells in young and aged CBA mice. Overall, our findings reveal a specific age-related hypoconnectivity of intra-cortical excitatory and inhibitory circuits from L4 in A1. The nature of the hypoconnectivity differed between males and females. Our results thus suggest that not all circuits are changing equally and that there are sex differences. Therefore, therapeutic interventions must take these laminar and sex-specific changes into account.

## Methods

All procedures were approved by Johns Hopkins University Institutional Animal Care and Use Committee

### Animals

Male and female CBA/CaJ mice (Jackson Laboratory #000654) were raised in 12-h light/12-h dark light cycle (N=12 adult group; N=14 aging group). Animals from P543 to P626 were used in the aging group (mean age P586), roughly matching ages from our in vivo study (Shilling-Scrivo et al., 2021). Young adult mice from P47 to P72 (mean age P62) were used as the controls.

### Slice Preparation

Mice were deeply anesthetized with isoflurane (Halocarbon). A block of brain containing A1 and the medial geniculate nucleus (MGN) was removed, and slices (400 μm thick) were cut on a vibrating microtome (Leica) in ice-cold artificial cerebrospinal fluid (ACSF) containing (in mM) 130 NaCl, 3 KCl, 1.25 KH_2_PO_4_, 20 NaHCO_3_, 10 glucose, 1.3 MgSO_4_, and 2.5 CaCl_2_ (pH 7.35–7.4, in 95% O_2_/5% CO_2_). Slices were cut ∼15 degrees from the horizontal plane to maintain the tonotopic organization in the slice (Zhao et al., 2009; Meng et al., 2015, 2017a; Meng et al., 2017b). Left hemisphere slices were incubated for 1 h in ACSF at 30°C and then kept at room temperature. For recording, slices were held in a chamber on a fixed-stage microscope (Olympus BX51) and superfused (2–4 ml/min) with high-Mg recording solution at room temperature to reduce spontaneous activity in the slice. The high-Mg recording solution contained (in mM) 124 NaCl, 5 KCl, 1.23 NaH_2_PO_4_, 26 NaHCO_3_, 10 glucose, 4 MgCl_2_, and 4 CaCl_2_. The location of the recording site in A1 was identified by landmarks (Cruikshank et al., 2002; Zhao et al., 2009; Meng et al., 2015).

### Electrophysiology

Whole-cell recordings were performed with a patch clamp amplifier (Multiclamp 700B, Molecular Devices) using pipettes with input resistance of 4–9 MΩ. Pyramidal cells targeted for recording were in an area of A1 overlying the rostral flexure of the hippocampus. Data acquisition was performed by National Instruments AD boards and custom software (Ephus) (Suter et al., 2010), written in Matlab (Mathworks) and adapted to our setup. Voltages were corrected for an estimated junction potential of 10 mV. Electrodes were filled with an internal solution containing (in mM): 115 cesium methanesulfonate (CsCH_3_SO_3_), 5 NaF, 10 EGTA, 10 HEPES, 15 CsCl, 3.5 MgATP, and 3 QX-314 (pH 7.25, 300 mOsm). Biocytin or neurobiotin (0.5%) was added to the electrode solution as needed. Series resistances were typically 20–25 MΩ.

### Laser-Scanning Photostimulation

LSPS was performed as described previously (Meng et al., 2015, 2017a; Meng et al., 2017b; Meng et al., 2020). Caged glutamate (0.8 mM *N*-(6-nitro-7-coumarylmethyl)-L-glutamate) (Muralidharan et al., 2016) was added to the high-divalent ACSF during recording. Laser stimulation (1 ms) was delivered through a 10x water immersion objective (Olympus). Laser power on the specimen was <25 mW and was held constant between recordings. For each map, an array of up to 30 × 30 sites with 30-μm spacing was stimulated once at 1 Hz in a pseudorandom order. This stimulation paradigm evokes an action potential at the stimulation sites with similar spatial resolution (about 100 μm) over cells in all cortical layers (Meng et al., 2015, 2017a; Meng et al., 2017b). Putative monosynaptic excitatory postsynaptic currents (EPSCs) in GABAergic interneurons were classified by the post-stimulation latency of the evoked current. Evoked currents with latencies of less than 10 ms are likely to be the result of direct activation of glutamate receptors on the patched cell. Evoked currents with latencies between 10 and 50 ms were classified as monosynaptic evoked EPSCs. The first peak amplitude and the charge (the area of EPSC in the counting window) were quantified for each synaptic response. Recordings were performed at room temperature and in high-Mg^2+^ solution to reduce the probability of polysynaptic inputs. Cells that did not show any large (> 100 pA) direct responses were excluded from the analysis, as these could be astrocytes. Excitatory and inhibitory inputs were recorded at −70 mV and 0 mV respectively.

### Statistics

Data were analyzed by custom software written in MATLAB. Cortical layer boundaries were identified by features in the bright-field image as described previously (Meng et al., 2015, 2017a; Meng et al., 2017b; Meng et al., 2020). Input area was calculated as the area within each layer that gave rise to PSCs. Integration distance referred to the distance that covered 80% of evoked PSCs along the rostro-caudal direction. Mean charge was the average charge of PSCs from each stimulus spot. And the mean peak amplitude was calculated by the average amplitude of PSCs. The balance of excitation and inhibition was calculated as the E/I ratio, which is based on the number of inputs and the average input strength (E_density_/I_density_ and E_charge_/I_charge_).

To determine the diversity of connection patterns in each group we calculated the spatial correlation of the binary connection maps in each group by calculating the pairwise cross-correlations (Meng et al., 2020). For pairwise correlation calculation, we set the area that has monosynaptic connections to 1 and the area without monosynaptic connections to zero. For each cell, we derive a 2-D matrix containing 0 and 1 which represent no connection spots and connection spots respectively. We then apply corrcoef function in matlab to obtain the correlation value between cells. The higher correlation suggests more similar connection patterns and lower correlation value indicates the heterogeneity of cortical circuits. The correlation coefficient is calculated based on the formula: *ρ*(*A,B*) = cov(*A,B*)/*σ*_*A*_*σ*_*B*_ Results are plotted as means ± SD unless otherwise indicated. Populations are compared with a rank sum test and Student’s t test.

## Results

To investigate the changes of intra-cortical circuits in A1 with aging, we performed LSPS, as described in our prior studies (Meng et al., 2015, 2017a; Meng et al., 2017b; Meng et al., 2020), in aged (older than 18 months) and young adult (P47-P72) CBA mice. Thalamocortical slices of A1 (Figure 1A) were cut and LSPS with caged glutamate was used to focally activate cortical neurons. Cell-attached patch recordings and whole-cell recordings were performed to test the photo-excitability of A1 neurons and the spatial connectivity of excitatory and inhibitory inputs to L2/3 neurons in A1 respectively (Figure 1A).

**Figure 1.**
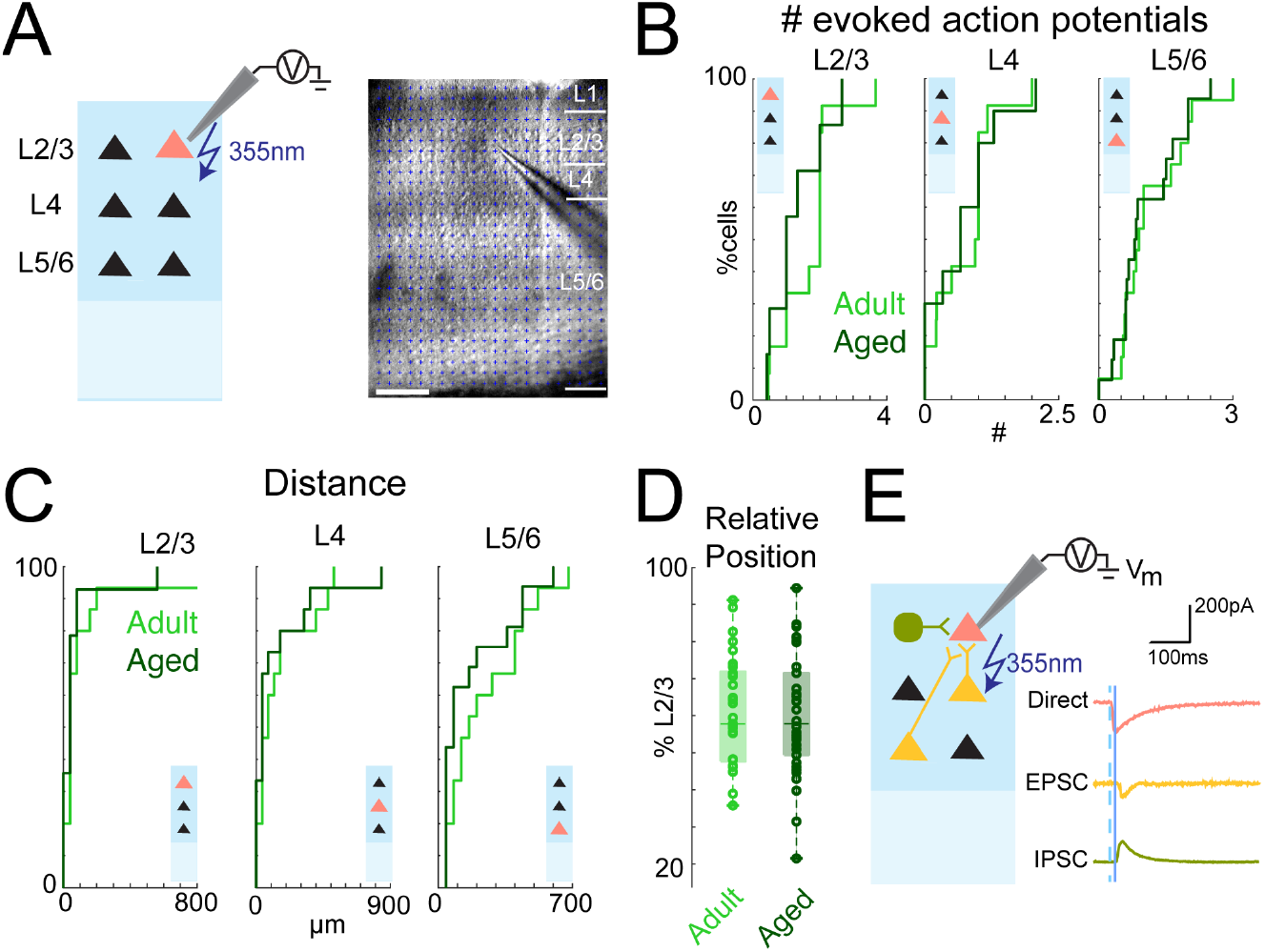
Photo-excitability of A1 Neurons Remains Unchanged During Aging. **A**, Left: schematic of LSPS for cell attached recording. Solid triangle represents recorded neuron. Right: Infrared image of brain slice with patch pipette on L2/3 neuron. Stimulation grid is indicated by blue dots. Layer boundaries are indicated by white bars on the right. The scale bar at the bottom left is 200 μm. **B-C**, Number of evoked action potentials (B) and effective stimulation distance (C) from cell-attached recordings of L2/3, L4 and L5/6 neurons. The numbers of evoked action potentials of neurons in all layers were similar and most spikes were evoked within 150 μm **D**, Relative position of recorded neurons within L2/3 relative to the borders of L4 and L1 (100%). Cells were sampled from a similar area in the middle of L2/3 in young adult and aged animals. **E**, Schematic of LSPS when recording evoked EPSCs and IPSCs. Whole-cell voltage clamp recordings were obtained from red cell at holding potentials of -70mV (EPSCs) or 0mV (IPSCs). If the presynaptic neuron (e.g. green or yellow cells) synapses on the recorded neuron, an PSC will be observed. Shown on the right are examplar patch-clamp recordings of direct response (red), EPSC (yellow), and IPSC (green), acquired at holding potentials of -70, -70, and 0 mV, respectively. Dashed blue line indicates time of photostimulation; solid blue line marks 8 ms post-stimulus, which is the minimal latency of synaptic responses.

### Aging does not alter photo-excitability of A1 neurons

To reliably compare the spatial connection pattern of cells in A1 using LSPS, we needed to confirm that the spatial resolution of LSPS was similar across ages. We performed cell-attached patch recordings with LSPS in cells from L2/3, L4 and L5/6 to test the ability of A1 neurons to fire action potentials in response to photoreleased glutamate (N=4, adult; N=3 aged). Short UV laser pulses (1 ms) were targeted to multiple stimulus locations to focally release glutamate and cause firing of action potentials. The grid of stimulation spots covered the entire A1, resulting in a high-resolution 2D photoactivation pattern for a given cell. We then counted numbers of evoked action potentials and measured the distance of effective stimulation sites. The numbers of evoked spikes were similar across ages (Figure 1B, p>0.05). Furthermore, the majority of action potentials were generated within 150 μm of the soma in both control and aged animals (Figure 1C, p>0.05). These findings suggest that the sensitivity of cells to photoreleased glutamate and thus the spatial resolution of LSPS remains unchanged with age

### L2/3 neurons in aged A1 show sex-specific reduction in intra-laminar excitatory connections

We next investigated if intracortical circuits to L2/3 pyramidal neurons change with aging [N=8 (4 females and 4 males), adult; N=11 (4 females and 7 males), aged]. To visualize the connection pattern impinging on L2/3 neurons, we combined LSPS with whole-cell recordings from A1 L2/3 cells. Recorded cells were located at similar laminar positions (Figure 1D, p>0.05) and held at -70 mV (E_GABA_) to isolate excitatory synaptic currents. Laser pulses were targeted to ∼900 distinct stimulus locations spanning all cortical layers around the recorded cell, and the resulting membrane currents were measured (Figure 1E). We observed large, short-latency (<10 ms) inward currents when the stimulation location was close to the recorded neuron due to direct activation of the cell body and the proximal dendrites. In contrast, post-synaptic currents caused by activation of presynaptic neurons were long-latency (>10ms) events (Meng et al., 2015, 2017a; Meng et al., 2017b; Meng et al., 2020).

We mapped 63 L2/3 cells in A1 (29 cells in 8 young adult animals and 34 cells in 11 aged animals). For each cell we identified stimulus locations that gave rise to an evoked EPSC and generated a binary input map. For each group, we aligned all maps to the soma position of the individual cells and averaged them (Figure 2A), resulting in a spatial connection probability map for excitatory inputs. These maps allowed us to identify cortical locations that over the population gave rise to inputs to L2/3 neurons. Qualitatively comparing the connection probability maps, we observed that maps from adult and aged animals were similar (Figure 2A).

**Figure 2.**
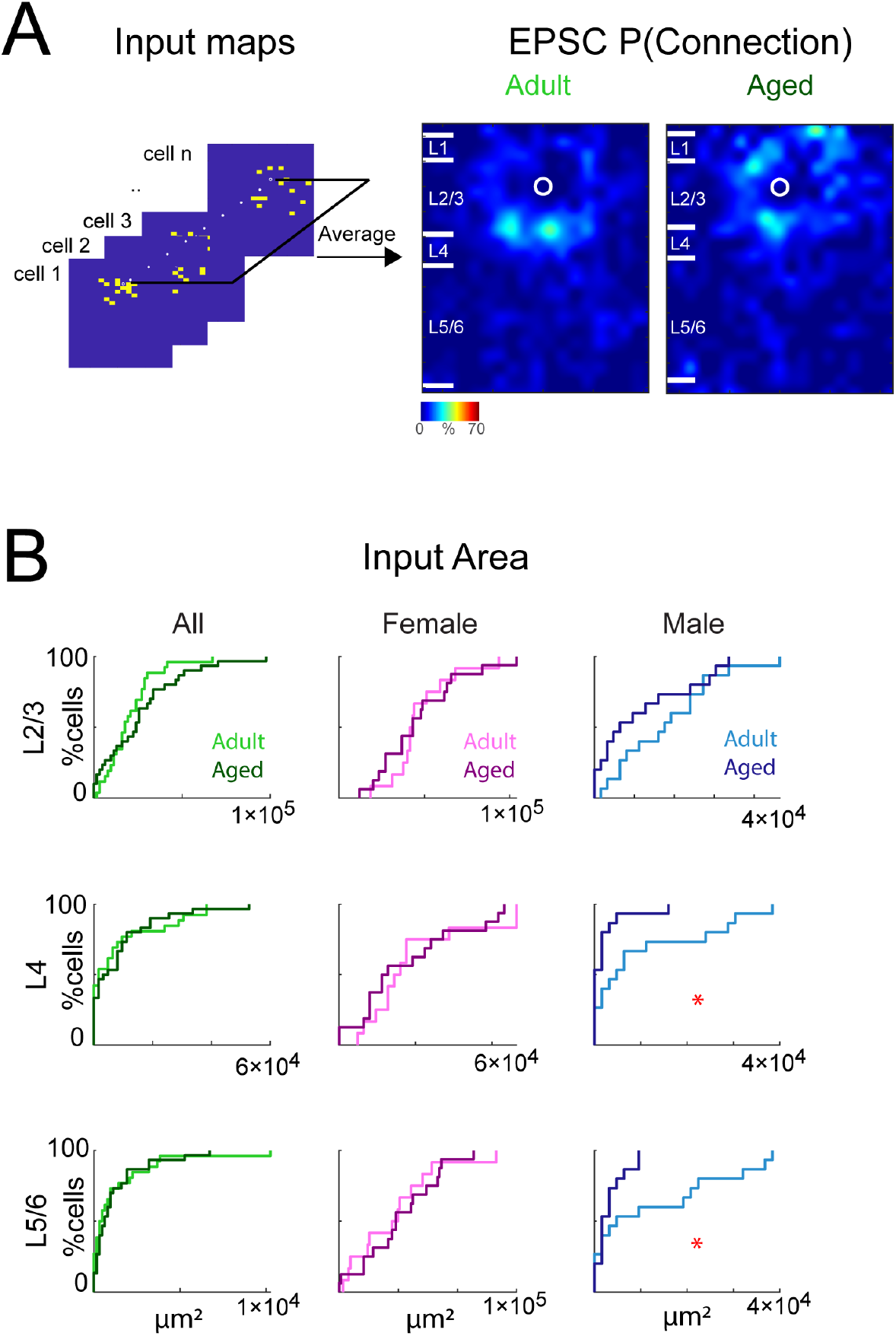
Sex-specific changes in excitatory connections to A1 L2/3 neurons in aged male mice. **A**, Left, Schematic illustration of how the connection probability map is calculated. Connection maps from recorded neurons are aligned to the soma location and averaged to create the connection probability map. Right: Maps of connection probability for excitatory connections in young adult (left) and aged (right) animals. Soma location is indicated by the white circle. Connection probability is encoded according to the pseudocolor scale. White horizontal lines indicate averaged laminar borders and are 100 μm long. **B**, Left, distributions of the area of excitatory input originating from L2/3 (top), L4 (middle), L5/6 (bottom) of young adult (light green) or aged (dark green) animals. Middle, distributions of the area of excitatory input originating from L2/3 (top), L4 (middle), L5/6 (bottom) of adult female (light pink) or aged female (dark pink) animals. Right, distributions of the area of excitatory input originating from L2/3 (top), L4 (middle), L5/6 (bottom) of young adult male (light blue) or aged male (dark blue) animals. *, p<0.05.

To quantify the amount of inputs from each layer, we calculated the laminar changes of the connection properties of each cell as in previous studies (Meng et al., 2015, 2017a; Meng et al., 2017b; Meng et al., 2020). We first identified layer boundaries in differential interference contrast (DIC) images. We next quantified the amount of convergence from each layer to L2/3. We calculated the total area within each layer from which EPSCs could be evoked. A comparison of input areas indicates that the amount of intralaminar excitatory input from within L2/3 and from L4 and L5/6 is similar in aging and adult mice (Figure 2B). In vivo imaging has revealed sex-specific differences between male and female mice with male mice showing a stronger effect of aging on sound-evoked responses and activity correlations (Shilling-Scrivo et al., 2021). We thus separately compared cells from male and female mice. We found that aging decreased excitatory inputs from L4 in male but not female mice (Figure 2B, median: 4.800×10^3^ vs 9.998×10^2^ p_ranksum_<0.028, mean 1.045×10^4^ vs. 2.027×10^3^ p_t-test_<0.025), with a trend towards decreased input evident also for inputs from L5/6 median: 4.800×10^3^ vs 1.600×10^3^ p_ranksum_ <0.222, mean 1.269×10^4^ vs. 3.200×10^3^ p_t-test_ <0.018). Thus, aging seems to affect the amount of ascending excitatory circuits to L2/3 neurons in male mice.

Our areal measurement accounts for both changes in the input distribution within a layer, e.g., from L2 to L3, as well as changes in the orthogonal direction. Our thalamocortical slices preserve the macroscale rostro-caudally oriented tonotopic map in the slice plane. Therefore, the spatial extent of the input distribution along the rostro-caudal axis is a proxy for the integration along the tonotopic axis. To probe if the reduction of L4 inputs occurred along the tonotopic axis, we next calculated the distance that includes 80% of the evoked EPSCs. We find that this intralaminar integration distance for inputs originating in L4 is reduced in aged male mice (Figure 3A, median: 280 vs 0 p_ranksum_ <0.043, mean 225 vs. 70.588 p_t-test_ <0.028), indicating that the same number of input originated from a smaller area of thalamorecipient layer in old male mice.

**Figure 3.**
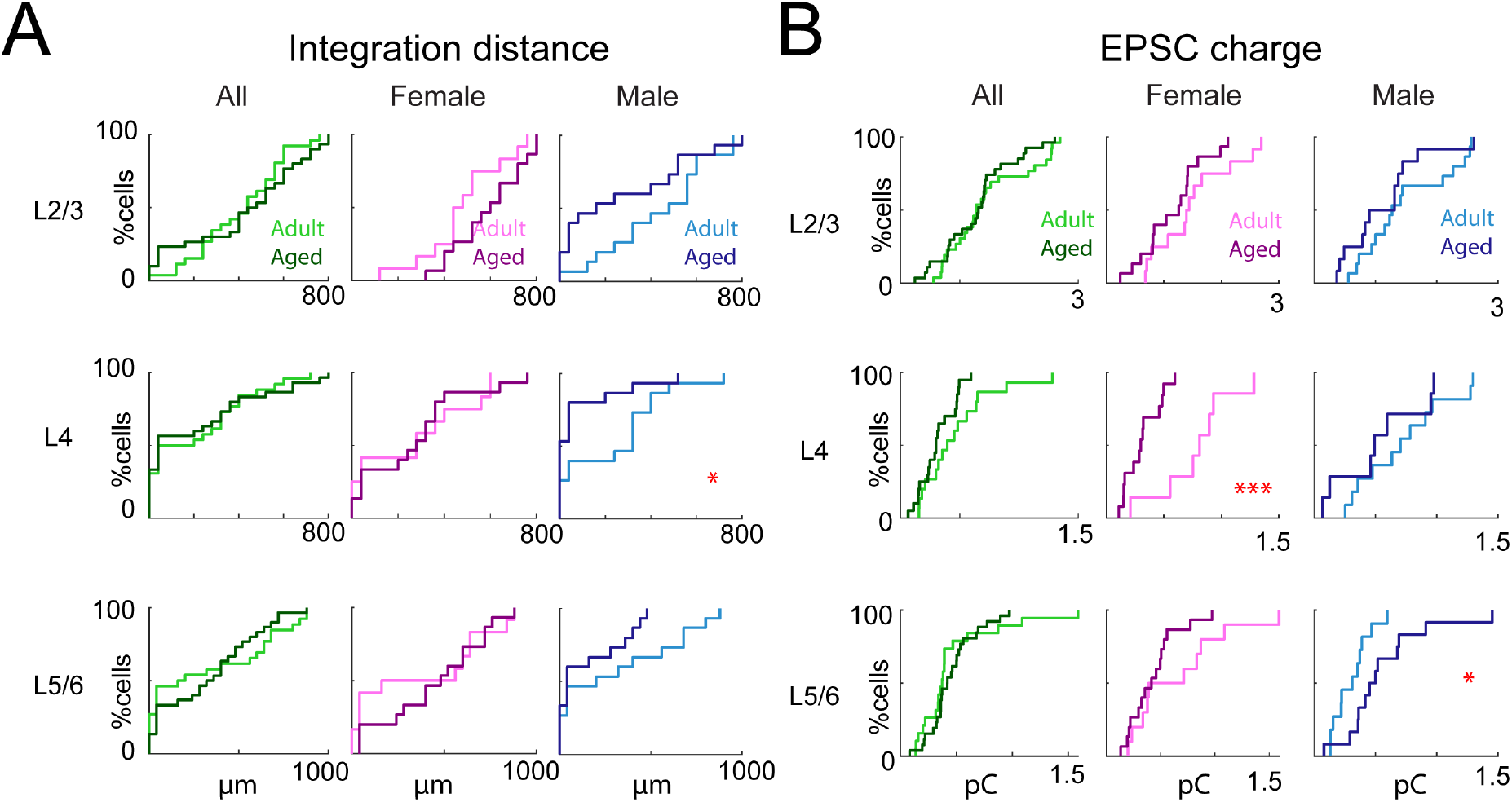
Sex-specific changes of connection strength to A1 L2/3 neurons in aged mice. **A**, Left, Integration distance to each L2/3 cell originating from L2/3 (top), L4 (middle), and L5/6 (bottom) of young adult or aged animals. Middle, Integration distance to each L2/3 cell originating from L2/3 (top), L4 (middle), and L5/6 (bottom) of adult female or aged female animals. Right, Integration distance to each L2/3 cell originating from L2/3 (top), L4 (middle), and L5/6 (bottom) of adult male or aged male animals. *, p<0.05. **B**, Left, distributions of mean EPSC charge of inputs originating from L2/3 (top), L4 (middle), and L5/6 (bottom) of young adult or aged animals; Middle, distribution of mean charge of inputs to each L2/3 cell originating from L2/3 (top), L4 (middle), and L5/6 (bottom) of adult female or aged female animals. ***, p<0.001. Right, distribution of mean charge of inputs to each L2/3 cell originating from L2/3 (top), L4 (middle), and L5/6 (bottom) of adult male or aged male animals. *, p<0.05.

Functional circuit changes can occur through alteration of connection probabilities but also through changes of connection strength. Given that many synaptic proteins change with aging, we next tested if connection strength was altered. We measured the average size (transferred charge) of the evoked EPSCs and found that connections from L4 in old females showed lower synaptic strength (median: 1.618 vs 0.596 p_ranksum_ <0.005, mean 1.554 vs. 0.625 p_t-test_ <0.0004), while L5/6 inputs in old males were strengthened (Figure 3B, median: 0.312 vs 0.477 p_ranksum_ <0.034, mean 0.306 vs. 0.560 p_t-test_ <0.038).

We next computed the fraction of inputs originating from each layer and found that cells from aging female and male mice received a similar fraction of inputs from all layers as young mice, with a trend to a reduced contribution from L4 in old male mice (Figure 4). These results indicate that L2/3 neurons in aged male mice tend to receive fewer interlaminar inputs especially from thalamorecipient L4.

**Figure 4.**
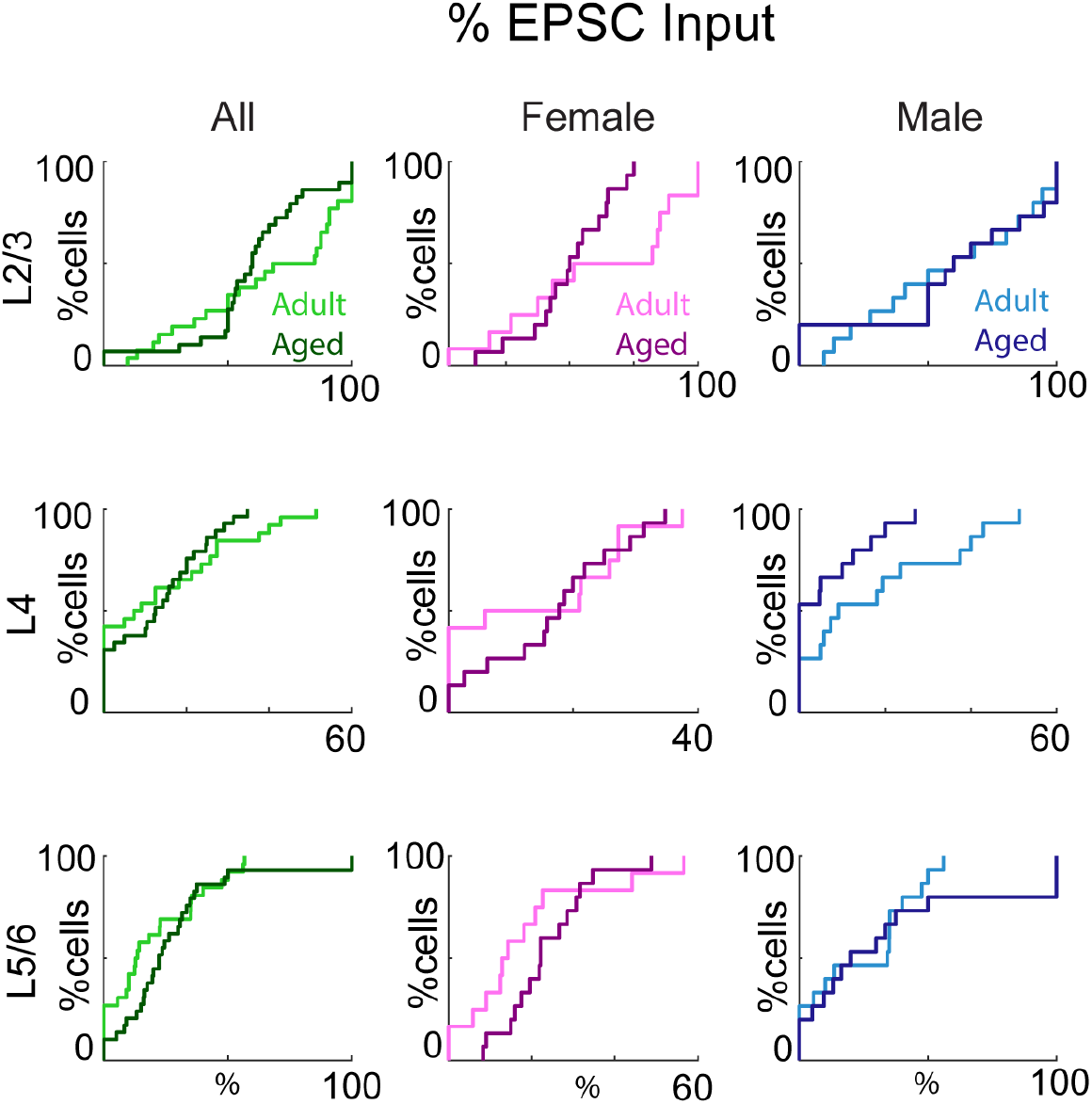
Percentage of excitatory inputs originating from each layer is similar between young and aged animals. Left, distributions of fraction of total excitatory input originating from L2/3 (top), L4 (middle), and L5/6 (bottom) of young adult or aged animals. Middle, distributions of fraction of total excitatory input to each L2/3 cell originating from L2/3 (top), L4 (middle), and L5/6 (bottom) of adult female or aged female animals. Right, distributions of fraction of total excitatory input to each L2/3 cell originating from L2/3 (top), L4 (middle), and L5/6 (bottom) of adult male or aged male animals.

Thus, these findings suggest that aged males have fewer inputs from L4 but stronger inputs from L5/6 while aged females show an unchanged connection pattern with weakened L4 inputs. Together, these data show a reduced contribution of excitatory L4 input to L2/3 in both males and females, but the mechanism of this reduction differed between males and females. Moreover, aged males show strengthened connection from L5/6. Thus, the circuit changes in A1 with aging vary by sex.

### L2/3 neurons in aged male A1 show reduced intralaminar inhibitory connections

We next investigated inhibitory circuits by holding cells at 0 mV and performing LSPS. We recorded evoked IPSCs and derived inhibitory input maps. Averaging these maps yielded inhibitory connection probability maps for both groups. In contrast to the excitatory maps, the connection pattern of inhibitory circuits showed few qualitative differences (Figure 5A). Laminar analysis quantitatively supported these observations. L2/3 neurons from young and old mice received a similar amount of inhibitory input from each layer. However, we observed significant decrease in the inhibitory connections from L4 in male but not female mice. (Figure 5B, median: 1.440×10^4^ vs 4.800×10^3^ p_ranksum_ <0.005, mean 1.497×10^4^ vs. 6.080×10^3^ p_t-test_ <0.004). Old male mice also showed a reduction in inhibitory inputs from L2/3 (median: 6.240×10^4^ vs 4.800×10^4^ p_ranksum_ <0.014, mean 7.966×10^4^ vs. 4.373×10^4^ p_t-test_ <0.014). However, the integration distance from all layers remains the same between young and aged mice, suggesting that the inputs are originating from similar area (Figure 6A). Together, our observations indicate that aging causes a reduction of inhibitory connections to L2/3 neurons in male mice.

**Figure 5.**
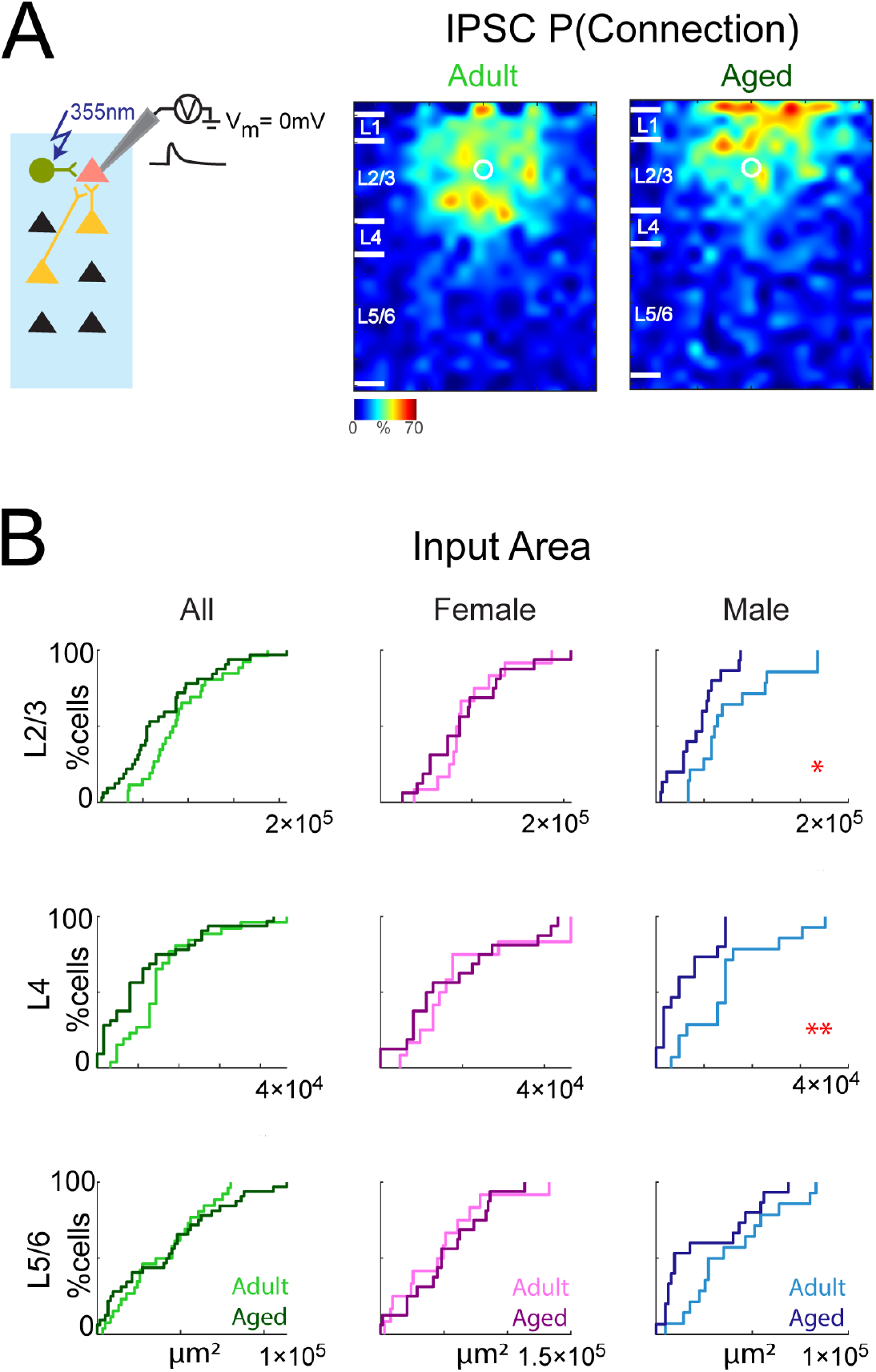
Sex-specific changes in inhibitory connections to A1 L2/3 neurons in aged male mice. **A**, Left, schematic of recording IPSCs. Whole-cell voltage clamp recordings at holding potentials of 0 mV. If the presynaptic neuron synapses onto the recorded neuron, an IPSC (black trace) will be observed. Right: Average maps of connection probability for inhibitory connections in young adult (left) and aged (right) animals. Maps are aligned to the soma location. Connection probability is encoded according to the pseudocolor scale. White horizontal lines indicate averaged laminar borders and are 100 μm long. **B**, Left, distributions of the area of inhibitory input originating from L2/3 (top), L4 (middle), L5/6 (bottom) of young adult (light green) or aged (dark green) animals. Middle, distributions of the area of inhibitory input originating from L2/3 (top), L4 (middle), L5/6 (bottom) of adult female (light pink) or aged female (dark pink) animals. Right, distributions of the area of inhibitory input originating from L2/3 (top), L4 (middle), L5/6 (bottom) of young adult male (light blue) or aged male (dark blue) animals. *, p<0.05; **, p<0.01.

**Figure 6.**
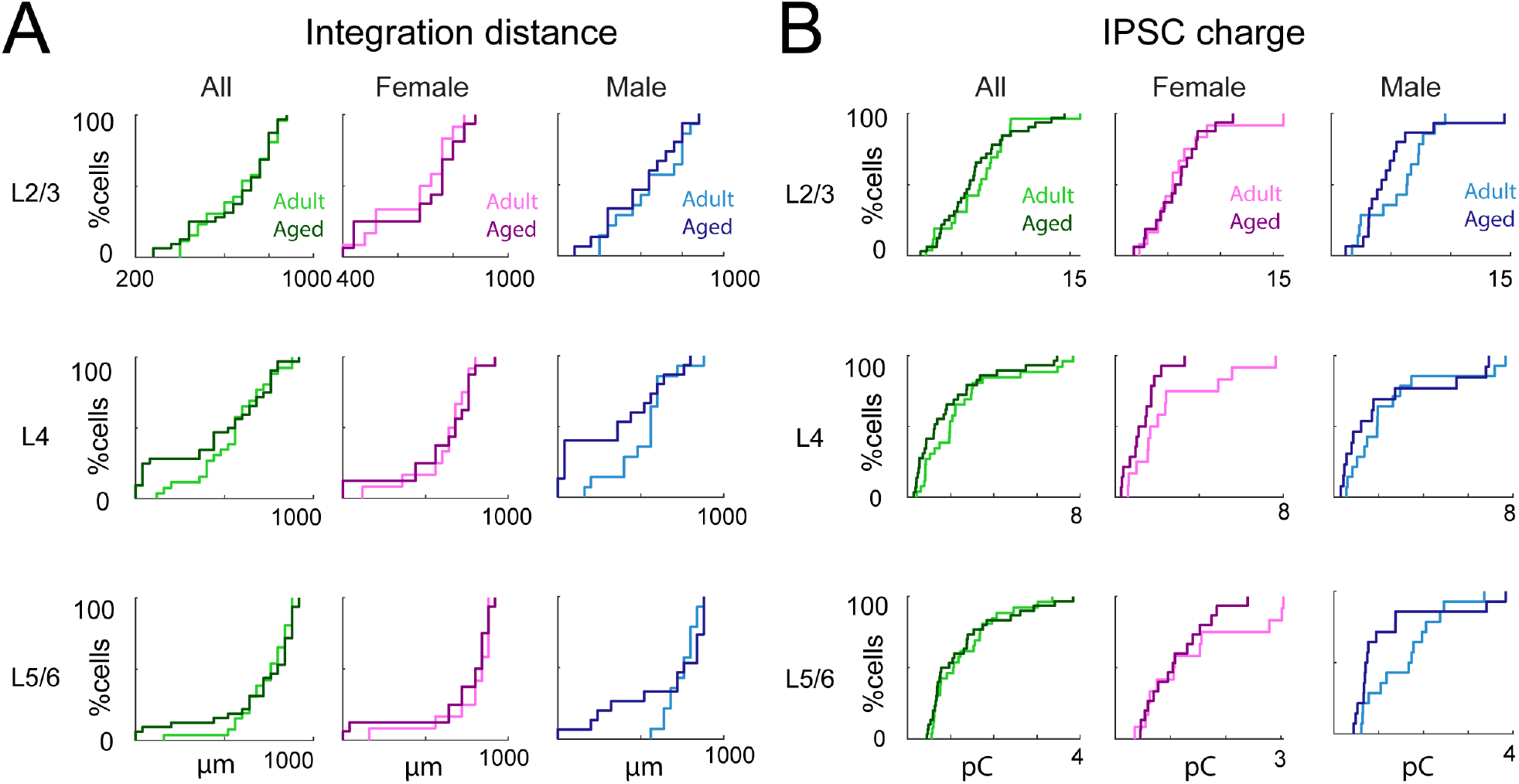
Inhibitory connection strength remain unchanged between young and aged animals. **A**, Left, Integration distance to each L2/3 cell originating from L2/3 (top), L4 (middle), and L5/6 (bottom) of young adult or aged animals. Middle, Integration distance to each L2/3 cell originating from L2/3 (top), L4 (middle), and L5/6 (bottom) of adult female or aged female animals. Right, Integration distance to each L2/3 cell originating from L2/3 (top), L4 (middle), and L5/6 (bottom) of adult male or aged male animals. *, p<0.05. **B**, Left, distributions of mean EPSC charge of inputs originating from L2/3 (top), L4 (middle), and L5/6 (bottom) of young adult or aged animals; Middle, distribution of mean charge of inputs to each L2/3 cell originating from L2/3 (top), L4 (middle), and L5/6 (bottom) of adult female or aged female animals. Right, distribution of mean charge of inputs to each L2/3 cell originating from L2/3 (top), L4 (middle), and L5/6 (bottom) of adult male or aged male animals.

Previous studies showed that the expression of rate-limiting enzymes, GAD65 and GAD67, which catalyze the conversion of glutamate to GABA, are decreased in aging in both IC and A1 (Milbrandt et al., 2000; Burianova et al., 2009), suggesting that strength of inhibition could be altered in aged mice. To test this hypothesis, we calculated the average charge of IPSCs. We found that IPSCs evoked from all layers had similar transferred charge independent of sex (Figure 6B).

We next computed the relative amount of inhibitory input from each layer and found that this was unchanged with age (Figure 7), indicating that inhibitory inputs from all layers were reduced in male mice with aging. Thus, our findings showed a reduction in the numbers of inhibitory inputs to L2/3 cells in old male mice.

**Figure 7.**
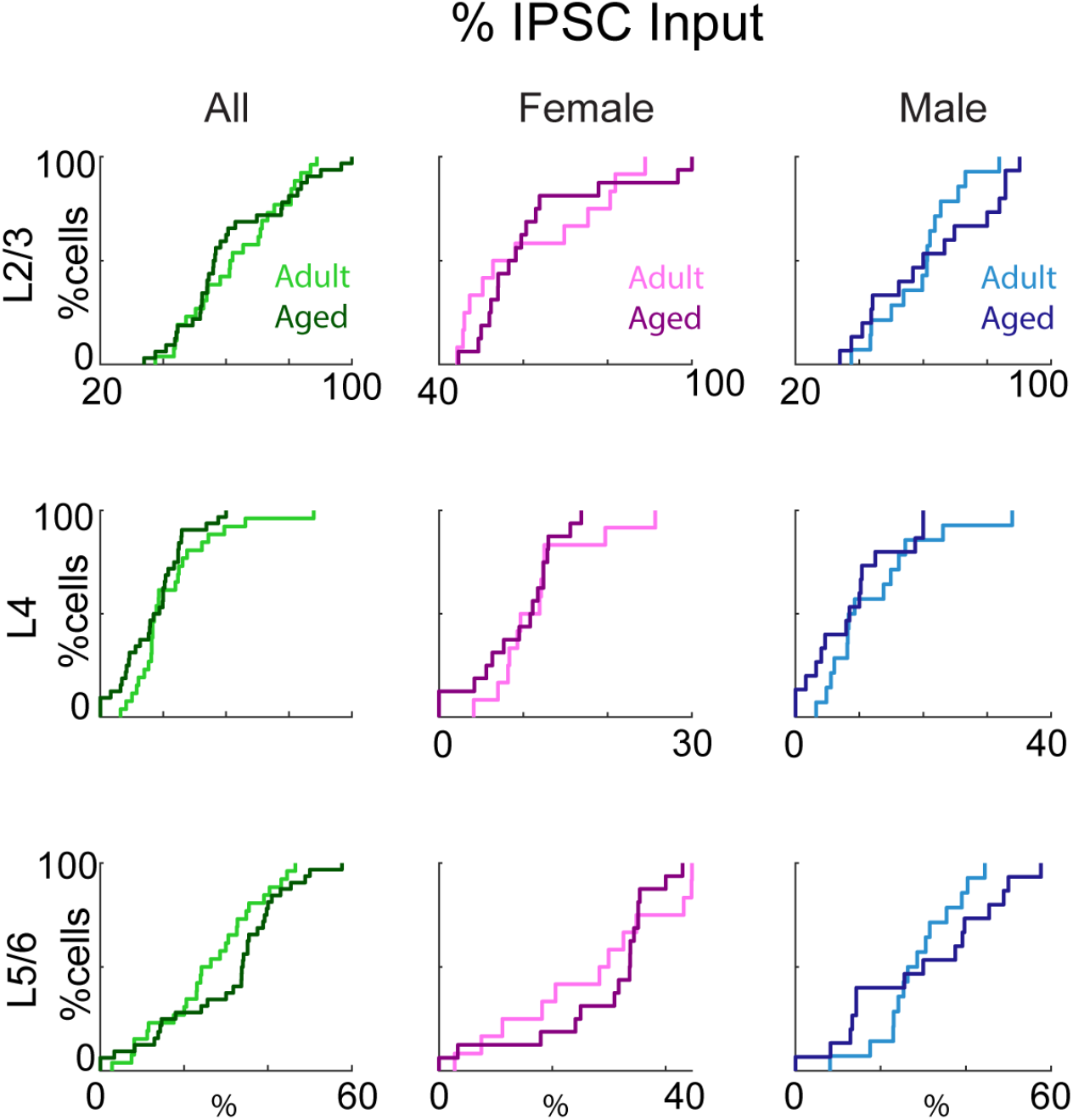
Percentage of inhibitory inputs originating from each layer is similar between young and aged animals. Left, distributions of fraction of total inhibitory input originating from L2/3 (top), L4 (middle), and L5/6 (bottom) of young adult or aged animals; Middle, distributions of fraction of total inhibitory input to each L2/3 cell originating from L2/3 (top), L4 (middle), and L5/6 (bottom) of adult female or aged female animals. Right, distributions of fraction of total inhibitory input to each L2/3 cell originating from L2/3 (top), L4 (middle), and L5/6 (bottom) of adult male or aged male animals.

### Excitation/inhibition balance in male mice is altered by aging

The concomitant occurrence of synaptic excitation and inhibition maintains balance between excitation and inhibition. In all sensory cortices, excitation and inhibition wax and wane together when responding to sensory stimuli (Anderson et al., 2000; Swadlow, 2003; Wehr and Zador, 2003; Tan et al., 2004; Wilent and Contreras, 2005), and the balance of these two opposing forces is critical for proper cortical function (Sohal and Rubenstein, 2019). Given that the E/I ratio changes and is crucial during development (Hensch and Fagiolini, 2005; Chu and Anderson, 2015; Hu et al., 2017), we asked whether E/I balance also changes during the aging process. We calculated the E/I balance of the inputs from each layer based on the density and the charge of PSCs as in our prior studies (Meng et al., 2015). We found that the E/I ratio based on density was reduced in aged males for inputs originating in L5/6 but not in L4 (Figure 8A, L5/6 median: 0.231 vs 0.160 p_ranksum_ <0.326, mean 0.455 vs. 0.135 p_t-test_ <0.0323; L4 median: 0.444 vs 0.111 p_ranksum_ < 0.162; Mean 0.505 vs 0.256 p_t-test_ < 0.092). However, E/I ratios based on charge (Figure 8B) were unchanged in aged animals. These results indicate that as the relative number of intralaminar excitatory inputs to L2/3 neurons decreased in old male mice, so did inhibitory inputs but to a larger amount.

**Figure 8.**
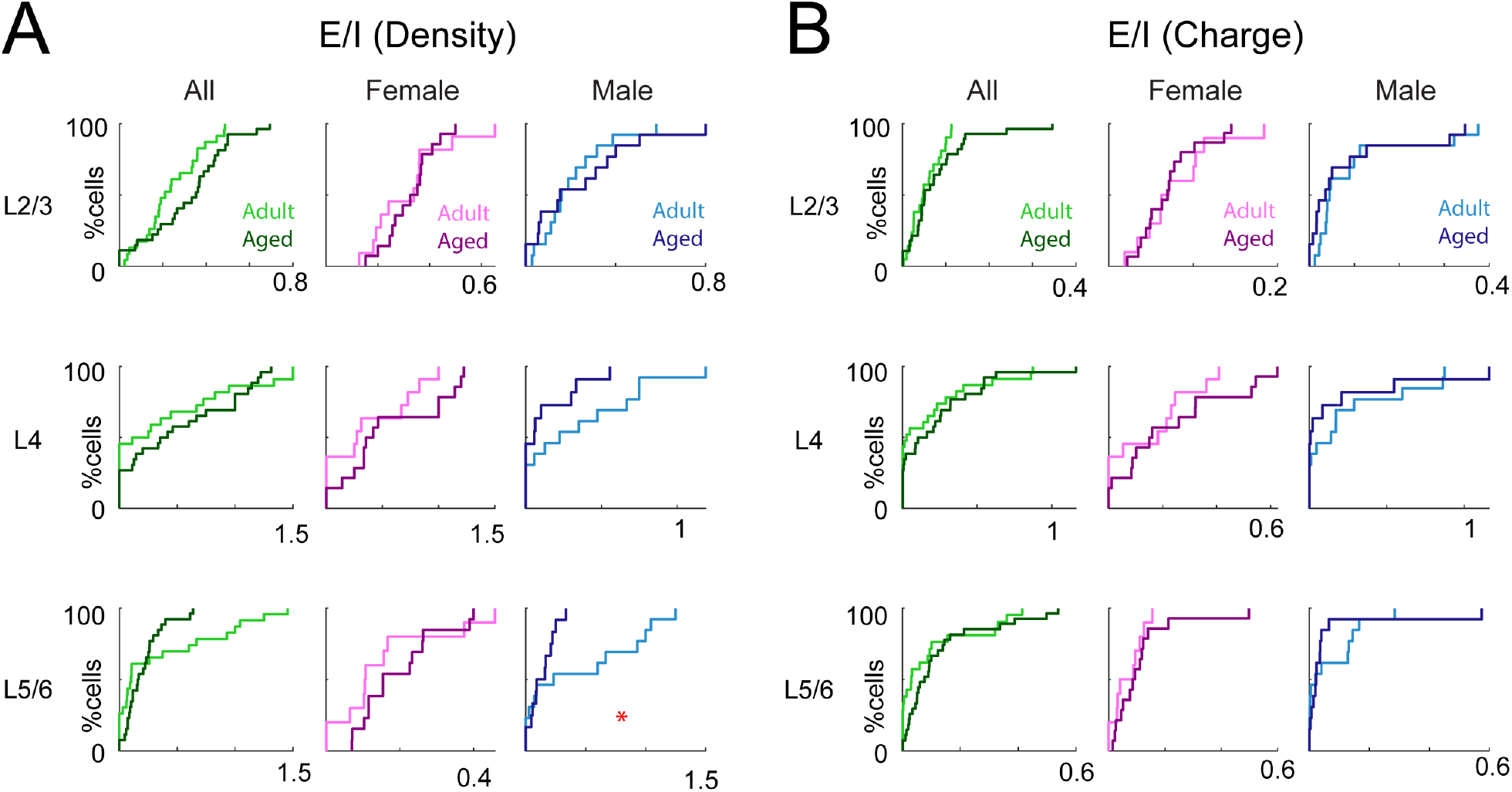
Male aged animals show altered E/I ratio. **A**, Left, distributions of E/I based on input numbers (density) from L2/3 (top), L4 (middle), and L5/6 (bottom) of young adult or aged animals. Middle, distributions of E/I based on input numbers (density) to each L2/3 cell originating from L2/3 (top), L4 (middle), and L5/6 (bottom) of adult female or aged female animals. Right, distributions of E/I based on input numbers (density) from L2/3 (top), L4 (middle), and L5/6 (bottom) of adult male or aged male animals. *, p<0.05. **B**, Left, distributions of E/I based on transferred charge originating from L2/3 (top), L4 (middle), and L5/6 (bottom) of young adult or aged animals; Middle, distributions of E/I based on transferred charge to each L2/3 cell originating from L2/3 (top), L4 (middle), and L5/6 (bottom) of adult female or aged female animals. Right, distributions of E/I based on transferred charge to each L2/3 cell originating from L2/3 (top), L4 (middle), and L5/6 (bottom) of adult male or aged male animals.

### Decreased circuit similarity in aged mice

L2/3 cells in adult A1 show heterogeneity in their functional interlaminar circuits (Meng et al., 2017a). This heterogeneity emerges over development, coinciding with a decrease in functional activity correlations (Meng et al., 2020).

To examine whether aging alters the circuit heterogeneity of L2/3 cells, we calculated the correlation between individual connection maps from all recorded cells (Figure 9A). We aligned the connection maps of recorded neurons within each group to their soma positions and compared the connection patterns between cells. This analysis showed that the correlation of excitatory and inhibitory circuits decreased in old mice due a decrease of similarity in L4 (Figure 9B, C). These effects are most pronounced in male mice consistent with the reduction of inputs from L4 in aged male mice. In aged female mice, inhibitory inputs from within L2/3 are more diverse.

**Figure 9.**
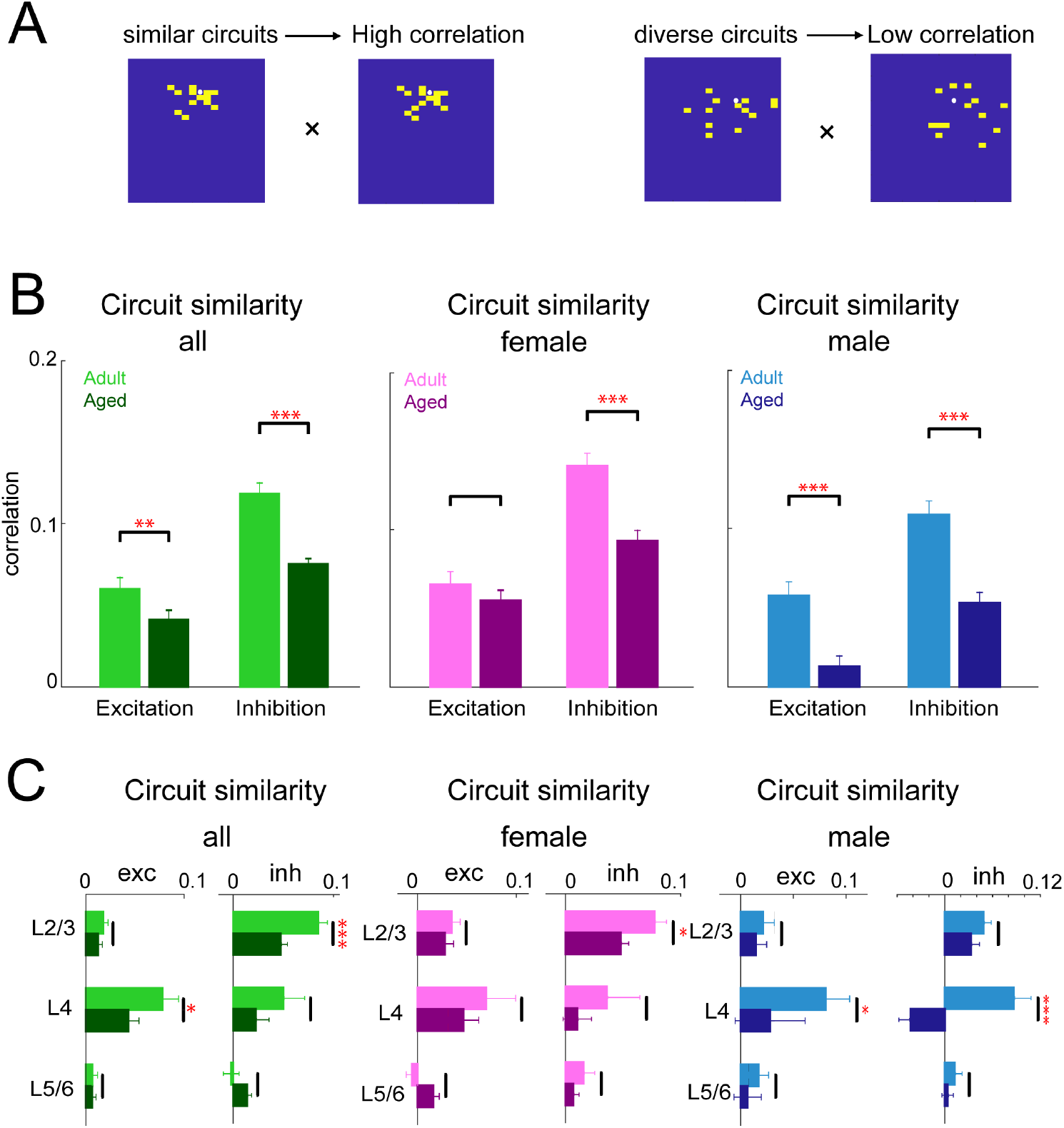
Decreased similarity of circuits in aged mice. **A**, Schematic illustration of how pairwise correlation between two maps is calculated. White circle represents recorded neuron. A yellow square represents a stimulus location that has monosynaptic connection to the recorded cell; a blue square represents a location with no connection to the recorded cell. For the pairwise correlation calculation, yellow and blue squares are assigned values of 1 and 0, respectively. **B**, Correlation of excitatory and inhibitory circuit patterns across all layers. Left, correlation of circuit patterns for all adult and aged animals. Middle, correlation of circuit patterns for adult female and aged female animals. Right, correlation of circuit patterns for adult female and aged female animals. **C**, Correlation of circuit patterns within L2/3 (left), L4 (middle) and L5/6 (right). Left, circuit similarity from different layers in adult and aged animals. Middle, circuit similarity from different layers in adult and aged female animals. Right, circuit similarity from different layers in adult and aged male animals. For all plots: *, p<0.05; **, p<0.01; ***, p<0.001.

Together, these results indicate that aging changes the spatial functional connection pattern of excitatory inputs to L2/3 neurons, increases the diversity of excitatory and inhibitory inputs from L4 in male mice and from L2/3 in female mice despite no changes in the amount of inputs.

## Discussion

Our results show a hypoconnectivity with reduced interlaminar excitatory and inhibitory connections to A1 L2/3 due to aging predominantly in male CBA mice. The circuit changes we observe are consistent with altered functional responses in A1 of aged CBA mice, which are more apparent in aged male mice (Shilling-Scrivo et al., 2021).

We observed a reduction of interlaminar excitatory inputs from L4 to L2/3 cells with age in male mice and a reduction in excitatory strength of these inputs in female mice. These results are consistent with the observation of reduced dendritic spine density in the aged brain (Dickstein et al., 2013; Hickmott and Dinse, 2013), and therefore suggest that these spines were innervated by L4 inputs. MGB cells, which provide input to L4, in aged animals show hyperexcitability to reduced tonic inhibition (Richardson et al., 2013), possibly increasing thalamocortical drive. Assuming a homeostatic framework, the decreased ascending interlaminar cortical connections to L2/3 could be a compensation mechanism for increased ascending drive from thalamocortical recipient layers or from higher order thalamic nuclei (Zhang and Bruno, 2019). We also observed a weakening of excitatory L4 connections in female mice. This is consistent with a loss of synaptic AMPA receptors and hypofunction of NMDA receptors as has been observed in the aged hippocampus and in other sensory areas (Gocel and Larson, 2013; Kumar et al., 2019).

Taken together, our results suggest that the aged A1 in CBA mice shows reduced excitation and that this reduction seems to affect ascending connections. Moreover, the relative decrease of excitatory L4 inputs we observe in male mice might contribute to the reduced bandwidth of aging L2/3 neurons (Shilling-Scrivo et al., 2021) as L4 neurons show broader tuning than L2/3 neurons (Winkowski and Kanold, 2013; Bowen et al., 2020).

Much focus has been placed on the changes in inhibition during aging (Chao and Knight, 1997; Caspary et al., 1999; Kok, 1999; Ling et al., 2005; Caspary et al., 2008; Stanley et al., 2012; Caspary et al., 2013; Rozycka and Liguz-Lecznar, 2017; Recanzone, 2018; Rogalla and Hildebrandt, 2020). Indeed, we did observe a reduction of functional inhibitory connections, but this was only present in male mice. The interneuron population is stable with aging (Ouellet and de Villers-Sidani, 2014; Burianova et al., 2015; Rogalla and Hildebrandt, 2020) and no age-related changes in the overall number, total binding, or affinity of GABA_A_ receptors have been observed (Heusner and Bosmann, 1981; Komiskey and MacFarlan, 1983; Reeves and Schweizer, 1983; Komiskey, 1987). Together with our data this points to a loss of inhibitory connections in male mice. However, prior studies did not separate animals by sex, thus it is possible, for example, that changes in cell number exist in aged male mice.

We found that the inhibitory input area originating from L4 is more concentrated in aged male animals and overall inhibition is more diverse between cells. Inhibition is critical for shaping tuning properties and information redundancy (Sillito, 1977, 1979; Kyriazi et al., 1996; Wang et al., 2000; Wang et al., 2002; Kurt et al., 2006; Razak and Fuzessery, 2009; Sun et al., 2010; Gaucher et al., 2013; Li et al., 2014; Zhou et al., 2014). A less dispersive input area from L4 might contribute to an abnormal tuning bandwidth (Turner et al., 2005; Shilling-Scrivo et al., 2021). We find that excitation from L5/6 is stronger in aged male animals. Inputs from L5/6 are thought to be important in gain control (Chambers et al., 2016), thus strengthening of these inputs could contribute to the increased population correlations observed in vivo (Shilling-Scrivo et al., 2021).

Our results show that the E/I balance remains stable with aging. This suggests a coordinated change of both excitation and inhibition. While an E/I imbalance has been found in the parietal cortex with age-related cognitive impairment, aged animals with normal cognition do not show an imbalance (Wong et al., 2006), consistent with our results.

Aging can alter the tonotopic map in A1 with aged animals showing a higher degree of scatter in frequency tuning and disorganized frequency distribution (de Villers-Sidani et al., 2010; Kamal et al., 2013). Meanwhile, aged animals show more diverse A1 receptive field possibly due to dysregulated plasticity in the aging cortex (Turner et al., 2005; Cisneros-Franco et al., 2018). In contrast, groups of cells in aged animals show increased functional correlations (Shilling-Scrivo et al., 2021). These results suggest a more heterogenous functional organization but more homogenous activation of aged A1 neurons. These results are consistent with our observation of increased diversity of intracortical circuits and changes in intracortical inhibitory circuits. Moreover, in addition to intracortical circuit changes functional changes might also reflect alterations in axonal bouton dynamics that are present in aged thalamocortical axons (Grillo et al., 2013).

We find a strong effect of sex on the age-related changes in A1 circuits in that changes are most pronounced in males but that distinct changes exist in females. Given that we see changes in the interlaminar PSC amplitudes in females that seem to mirror the changes in connections in males, we speculate that central aging in males might be accelerated compared to females (Henry, 2004). This effect of sex we reveal is consistent with in vivo studies that show that male CBA mice have much larger age-related increases in correlated activity than females (Shilling-Scrivo et al., 2021). These results strongly suggest that much more work needs to be done in humans and other model species to see if this extends beyond rodents. Indeed, already sex differences have been found in human cortical synaptic connections (Alonso-Nanclares et al., 2008). Moreover, given that we see distinct differences in the age-related changes between males and females our data suggest that therapeutic approaches might need to be tailored by sex.

Taken together, our findings delineate effects of aging on mouse primary auditory cortex on a microcircuit level. We demonstrate that aging has more dramatic effects of A1 in male CBA mice, resulting in a loss of interlaminar excitatory and inhibitory circuits in aged males. Our results suggest that the altered response properties to sound in aged male CBA A1 are due to the changes of both excitatory and inhibitory connections.

## Acknowledgments and Contributions

BX and POK conceived study. BX performed all experiments and analyzed data. JPYK provided reagents. POK, BX wrote the manuscript. Supported by NIH P01AG055365 (POK), NIH RO1DC009607 (POK), NIH R01GM056481 (JPYK).

